# Mitotic waves in frog egg extracts: Transition from phase waves to trigger waves

**DOI:** 10.1101/2024.01.18.576267

**Authors:** Owen Puls, Daniel Ruiz-Reynés, Franco Tavella, Minjun Jin, Yeonghoon Kim, Lendert Gelens, Qiong Yang

**Author notes:** contributed equally (Owen Puls, Daniel Ruiz-Reynés). contributed equally (Lendert Gelens, Qiong Yang).

## Abstract

Cyclin-dependent kinase 1 (Cdk1) activity rises and falls throughout the cell cycle, a cell-autonomous process known as mitotic oscillations. These oscillators can synchronize when spatially coupled, providing a crucial foundation for rapid synchronous divisions in large early embryos like *Drosophila* (*∼* 0.5 mm) and *Xenopus* (*∼* 1.2 mm). While diffusion alone cannot achieve such long-range coordination, recent studies have proposed two types of mitotic waves, phase and trigger waves, to explain the phenomena. How the waves establish over time for efficient spatial coordination remains unclear. Using *Xenopus laevis* egg extracts and a Cdk1 FRET sensor, we observe a transition from phase waves to a trigger wave regime in an initially homogeneous cytosol. Adding nuclei accelerates such transition. Moreover, the system transitions almost immediately to this regime when externally driven by metaphase-arrested extracts from the boundary. Employing computational modeling, we pinpoint how wave nature, including speed-period relation, depends on transient dynamics and oscillator properties, suggesting that phase waves appear transiently due to the time required for trigger waves to entrain the system and that spatial heterogeneity promotes entrainment. Therefore, we show that both waves belong to a single biological process capable of coordinating the cell cycle over long distances.

## INTRODUCTION

Cell division, one of the most important processes in biology, is regulated by a well-studied pacemaker oscillator centered on the cyclin B-Cdk1 complex, known as the mitotic clock^1,2^ (Fig. 1**a**). The mitotic cell cycle, specifically the DNA-replication-and-division cycle, undergoes a sequence of events in which cells replicate DNA and partition the copies into daughter cells such that each daughter receives precisely one copy of the genome^3^.

**Fig. 1.**
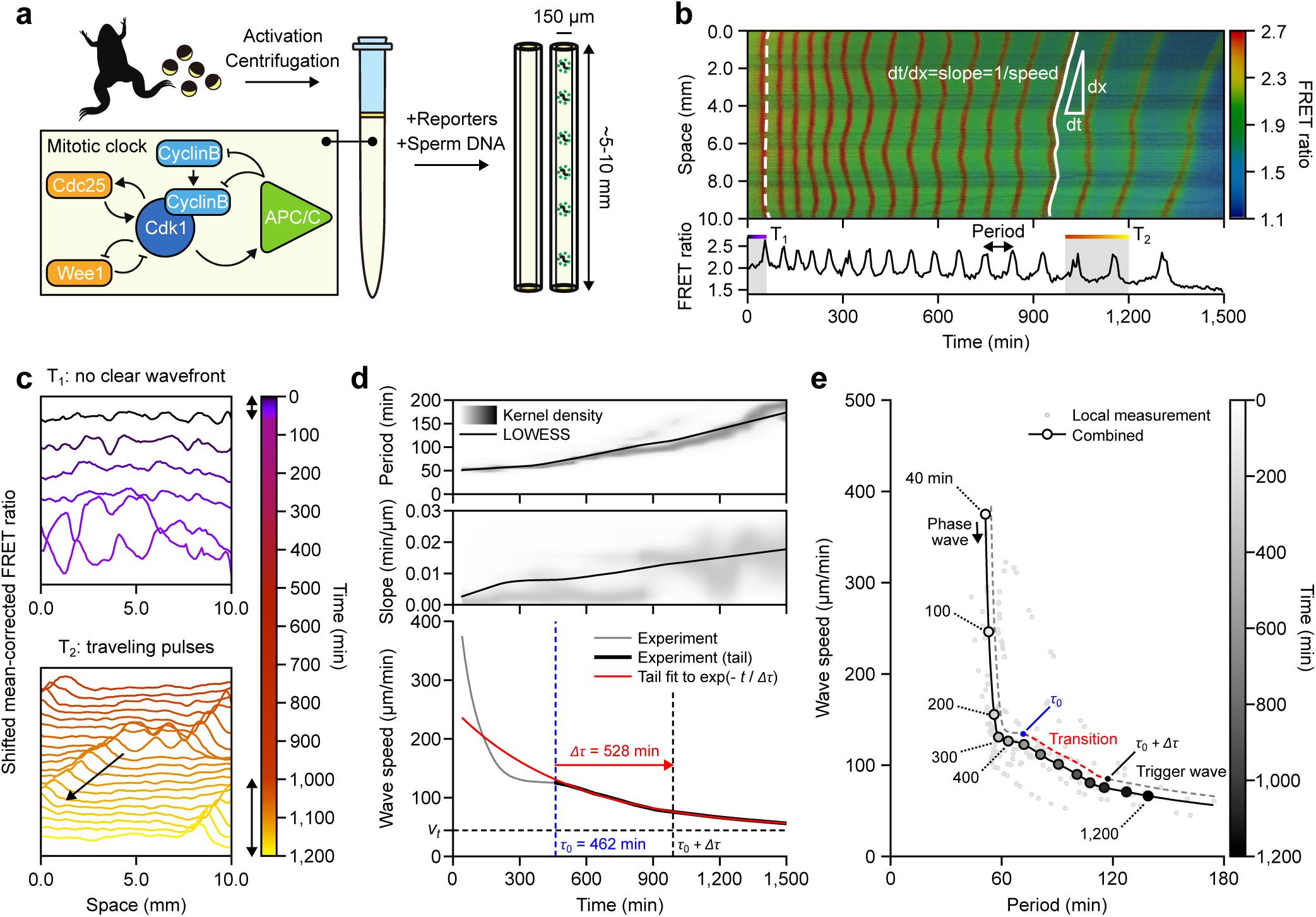
Time evolution of mitotic waves. **a** Schematic view of the experimental preparation of cycling egg extracts of *Xenopus laevis* supplied with Cdk1-FRET sensor, as a reporter, and loaded in Teflon tubes of *∼* 5-10 mm long and 150 µm of inner diameter. Before loading, extracts were added with *Xenopus* demembranated sperm DNA (+XS) or without (*−*XS, Control). Bottom-Left: Schematic representation of the regulatory network driving the mitotic oscillations. **b** Top: FRET ratio kymograph for a representative tube of 10 mm long loaded with extracts without sperm DNA (*−*XS). The top end of the tube is assigned to *x* = 10 mm. Color bar indicates FRET ratio values. For illustration purposes, two detected wavefronts are labeled in white lines (dashed for a wave at an early time and solid for a later-time wave). The slope (*dt/dx*) of these wavefronts describes the change in time with respect to traveling distance and is the inverse of speed. Bottom: FRET ratio time course recorded at the bottom end of the Teflon tube (*x* = 10 mm). The period is defined as the time interval between consecutive FRET ratio peaks. One early time region, *T*_1_, and one late time region, *T*_2_, are selected for further analysis in panel **c**. **c** Shifted mean-corrected FRET signal for two different time regions, *T*_1_ and *T*_2_, defined in B across the Teflon tube. Top: Early-time signal showing in-phase activation with no clear wavefront. Bottom: Late-time signal showing a traveling pulse crossing the tube over time, as indicated by the black arrow. Bidirectional arrows on the colorbar indicate *T*_1_ and *T*_2_ time regions. **d** Time evolution of the period (top), slope (middle), and wave speed (bottom). Both period and slope are measured locally for each wavefront locus. Their distributions are represented using the kernel density estimation (KDE), indicated with a grayscale colormap, and normalized at each time. Period and slope data are smoothed by locally weighted scatterplot smoothing (LOWESS) and their curves are shown with solid black lines. The wave speed is obtained as the inverse of the LOWESS estimation of the slope. The time point at which exponential decay begins (*τ*_0_, dashed blue line) is calculated via a moving horizon fitting (See Methods and Fig. S1 for the definition). The evolution of the speed after *τ*_0_ is fitted by an exponential function (solid red line) to calculate the entrainment time (Δ*τ*). The horizontal dashed black line indicates the resulting terminal velocity from the fit (*v_t_*) and the vertical dashed black line indicates the time point corresponding to *τ*_0_ + Δ*τ*. **e** Speed-period relationship. Local measurements of the period and wave speed are represented by gray dots (2% of all data points were shown). Combined speed-period relation (solid black line) is computed from the LOWESS estimations of the period and speed from **d**. Additional markers (open circles with grayscale fillings) are placed along this line to indicate multiples of 100 minutes (except for the first one which is shown for clarity). Grayscale color bar indicates time. The transition time points *τ*_0_ and *τ*_0_ + Δ*τ* are indicated on a dashed guideline with the transition time frame highlighted in red. Data in **d** and **e** are pooled from two independent *Xenopus* egg batches and three replicates each.

In the early embryogenesis of organisms such as *Xenopus* or *Drosophila*, cells initially proceed through a series of fast divisions^4,5^. These mitotic cycles lack many features of mature cells—e.g., gap phases, cell cycle checkpoints, and zygotic gene transcription—which only arise after the mid-blastula transition (MBT)^6^. For this reason, it is key for these cycles to remain roughly synchronized prior to MBT, even though some desynchronization is tolerated^7^. Throughout this process, mitotic events occur within minutes of each other. However, due to the large cell size in such embryos, diffusion alone remains far too slow to synchronize the system: such a process would take multiple hours, not minutes^8–12^.

Previous studies have identified waves of mitotic events, both *in vitro* and *in vivo*, which propagate at speeds sufficiently high to communicate across the lengths of the embryo^8,9^. Trigger waves (*∼* 40-60 µm/min), resulting from the coupling of diffusion and local dynamics^10,11^, were first shown to coordinate mitosis in *Xenopus*, using nuclear envelope breakdown (NEB) to illustrate their propagation after a few early, largely synchronous cycles^8^. Subsequent work revealed that the nucleus itself serves as the pacemaker for these waves, locally accelerating oscillations possibly by aggregating cell cycle regulators and thus driving waves^13,14^. This aggregating mechanism was later confirmed in individual oscillating microemulsions^15^. However, the classical trigger wave mechanism may not be the sole contributor to the fast wave propagation observed in the early embryos.

Like *Xenopus* and other metazoans, the fruit fly embryo undergoes a series of rapid, roughly synchronous (and in this case, syncytial) divisions post-fertilization^6^. However, *Drosophila* embryos display waves of mitotic completion that traverse the entirety of the embryo (hundreds of microns) in mere minutes at early stages, with speeds much faster (*∼* 100 µm/min) than what could be achieved by traditional trigger wave models^9^. Moreover, embryos exhibit distinct spatial dynamics that forgo the classical picture of a stable regime invading into and promoting a metastable regime^10,11^. Instead, spatial gradients in Cdk1 activity are largely preserved, while the overall levels are swept upwards^16^.

Vergassola *et al*. propose that a distinct phenomenon termed “sweep” waves (or “phase” waves in our terminology, as accepted in a subsequent review^12^) is responsible for the ultra-fast waves observed *in vivo*^12,16^. Phase waves appear to spread due to local phase gradients but are not actively spread by mutual interactions, in contrast with trigger waves, which do propagate through a coupling of diffusion and local reactions^8–12^. The authors suggest that a time-dependent sweeping-up of Cdk1 activity leads to wave-like behavior spreading at scales faster than trigger waves, consistent with phase waves. Interestingly, the authors observe that the period of the oscillator lengthens for late cycles (due to Chk1) and propose a potential link between phase and trigger waves through this mechanism^16^. More recently, the authors speculate that such a transition could also exist in *Xenopus* and called for direct measurements of Cdk1 activity to resolve this open question^17^.

In this work, we present direct evidence of mitotic waves in *Xenopus* using a FRET sensor that measures the ratio of activity between Cdk1 and its opposing phosphatase. We show that waves of Cdk1 activity can form spontaneously in the absence of nuclear pacemakers, sharing the fundamental nature of the classical chemical waves in a Belousov-Zhabotinsky (BZ) reaction-diffusion system. We investigate the time-dependent behavior of mitotic waves in *Xenopus* extracts, revealing a transition from phase-wave-like to trigger-wave-like patterns over time, and as a result, offer the connecting thread between these phenomena. We also probe the role of nuclei in wave propagation, showing that in addition to acting as pacemakers, nuclei accelerate the entrainment of the system to the trigger wave regime. Building on the findings in these experiments, we then propose a novel method for generating directed waves *in vitro*, which reinforces the notion of entrainment explicitly. In short, we use a reservoir of active Cdk1 to drive waves through the oscillating medium. Taken together, we offer a generalized picture of the interplay between phase and trigger waves and the role of heterogeneities in the spatial coordination of *Xenopus laevis*’ early embryogenesis.

## RESULTS

### Mitotic waves transition from fast phase waves to slower trigger waves

We leverage *in vitro* cell-free extracts to characterize mitotic waves in *Xenopus*. A schematic view of the experimental setup is presented in Fig. 1**a**. Cycling extracts are prepared following the protocol described in previous studies^18,19^. Instead of relying on downstream events such as NEB, we employ a FRET sensor to report the Cdk1 kinase activity, which allows us to directly visualize mitotic waves over time in *Xenopus*^15^. Cycling extracts supplemented with the Cdk1 FRET sensor are then loaded into *∼* 5-10 mm long Teflon-coated tubes, submerged under mineral oil, and imaged using time-lapse epifluorescent microscopy.

In a representative experiment (Fig. 1**b**; Mov. 1), the FRET signal is represented by a heatmap with cool colors corresponding to low Cdk1 activity, and warm colors to high activity. High activity regions can be clustered together via peak detection, allowing us to individualize wavefronts. Two wavefronts at different time regions are highlighted for comparison (Fig. 1**b**, top, white lines). Qualitatively, one can observe a difference between early time patterns that are largely synchronous and fast moving (Fig. 1**b**, top, dashed white line), and later time patterns which form linear fronts (Fig. 1**b**, top, solid white line). The explicit time evolution of the FRET signal is also depicted for a small slice (at position *x* = 10 mm; Fig. 1**b**, bottom). When plotting the FRET ratio spatial profile over consecutive frames for early and late cycles (Fig. 1**c**), we observe clear changes in spatial profiles over time. Early patterns (*T*_1_: 0-60 min) resemble phase waves: a roughly uniform upswing in activity, and preservation of local peaks and spatial gradients (Fig. 1**c**, top), sharing similar features as those reported in Hayden *et al*.^17^. Conversely, at late times (*T*_2_: 1000-1200 min), the system exhibits clearly linear, trigger-wave-like fronts, characterized by traveling pulses (Fig. 1**c**, bottom). This implies a transition from phase waves at early, fast cycles, to trigger waves at late, slower cycles.

To quantify this transition, we choose to measure the period and wave speed. The period is calculated as the peak-to-peak time between wavefronts (Fig. 1**b**, bottom; Fig.1**d**, top). Since the speed of early time patterns is often infinite at this resolution, we calculate the instantaneous derivative (slope, *dt/dx*) of the interpolated kymograph, using its reciprocal as an indicator of wave speed. In both cases, we estimate the kernel density (KDE) of the data over time. We observe that the LOWESS estimate for the period closely follows the peaks in density (Fig. 1**d**, top). However, this is not the case for the slope. The slope density shows low values at early times, high values at later times, and a mixture in between, suggesting a transition between different types of waves (Fig. 1**d**, middle). Wave speeds are obtained by inverting the LOWESS slope estimate, showing a monotonic decrease over time (Fig. 1**d**, bottom). The decrease in wave speed seems to follow a two-step process with an initial fast decay and a slower transition to a terminal speed (*v_t_*). We quantify this transition following a moving horizon fitting algorithm (see Methods). Briefly, we find the potential transition starting time point (*τ*_0_) that gives the best fit for exponential decay of the signal at late times (Fig. S1). This time point tells us when waves start to relax exponentially towards the trigger wave state of later times. From the fitted exponential function, we extract a relaxation time scale (Δ*τ*) and thus fully quantify how long the system takes to transition between each state. For this data, wave speeds start to decay exponentially after *τ*_0_ = 462 min (Fig. 1**d**, bottom, dashed blue line) with a relaxation time scale of Δ*τ* = 528 min (Fig. 1**d**, bottom, red arrow). Interestingly, despite changes in period and wave speed, the FRET ratio maximum activation rate (*dA/dt*), calculated as the largest time derivative of the FRET ratio per cycle, remains relatively constant for the duration of the experiment (Fig. S2).

When combined, our measurements reveal that the wave speed monotonically decreases as the cell cycle period lengthens (Fig. 1**e**). At early times (before *τ*_0_), the period is short, and the system exhibits phase waves at diversified speeds of 400-100 µm/min, which are much faster than trigger waves. However, these transients eventually die off as the system transitions to a regime quasi-dominated by trigger waves (100-50 µm/min). This speed-period relation also appears to confirm a sweep-to-trigger transition, reported in fly embryos upon genetic perturbations^17^, though in this case, we directly observed the transition as an inherent temporal evolution of the system, independent of external induction, thus bridging our understanding of mitotic waves between different model systems.

### A generic cell cycle model shows that transient dynamics explains the observed phase to trigger wave transition

The observed transition from fast phase waves to slow trigger waves could be a result of two time-dependent factors. On the one hand, we observed period lengthening, which suggests a potential time dependence in the intrinsic biochemical properties of the oscillator. On the other hand, if we consider trigger waves as the attractor state of a dynamical system, the transition may imply a relaxation towards this state, a time-evolving process that necessitates a finite amount of time for a transient state (from a wide variety of initial conditions) to establish into a stable solution (attractor).

We turn to mathematical modeling to quantify the effect of these two hypothetical contributions to the wave dynamics we observed. We use a cell-cycle model introduced by Yang and Ferrell^20^ and then later extended by Chang and Ferrell^8^ to describe mitotic waves. The model describes the time evolution of active (denoted as *a*) and total cyclin B-Cdk1 (denoted as *c*) concentration (Eqs. (1) and (2)). Cyclin B is synthesized at a rate *k_s_*and then rapidly binds to Cdk1. The activity of the cyclin B-Cdk1 complex is regulated by phosphatase Cdc25, which activates the complex by dephosphorylation, and kinases Wee1 and Myt1, which deactivate the complex by phosphorylation. Finally, high Cdk1 activity leads to the activation of anaphase-promoting complex/cyclosome (APC/C), which targets cyclin B for degradation (Fig. 1**a**). The reaction rates are described by ultrasensitive response curves dependent on the cyclin B-Cdk1 activity and parameterized based on experiments^20^. Diffusion is incorporated into the model to simulate spatially extended dynamics.

To replicate the observed period lengthening, we introduce an explicit time dependence to the mitotic regulatory network. Adopting a methodology similar to Rombouts and Gelens^21,22^, we explore how the model parameters influence the period and activation rate (*da/dt*), which we use to compare to their experimentally observable counterparts (period and maximum activation rate *dA/dt*, extracted from the FRET signal time traces, respectively). We further modify the model equations with dimensionless factors *α*, *β*, *η* to examine the significance of the key network components (Fig. 2**a**). Factor *α* scales the activation and inactivation rates of Cdk1, which constitute the bistable switch at the mitotic network core. Numerical simulation shows that the period remains almost unaffected by *α* (Figs. 2**a** and S3, left). Factor *β* controls how fast cyclin B is synthesized and degraded, thus being fundamental for driving oscillations through the bistable trigger and negative feedback. Scaling *β* leads to a modulation of the period (interphase lengthening) and mostly unchanged activation rate (Figs. 2**a** and S3, middle). Finally, factor *η* corresponds to a global time scaling, thus affecting every observable (Figs. 2**a** and S3, right). Out of the three modifications, changing *β* produces the most similar behavior to our experimental observations: the period lengthens without a significant change in activation rate. Therefore, by dialing the bistable trigger over time (decreasing *β*), we obtain a phenomenological model that reproduces the observed cell cycle behaviors.

**Fig. 2.**
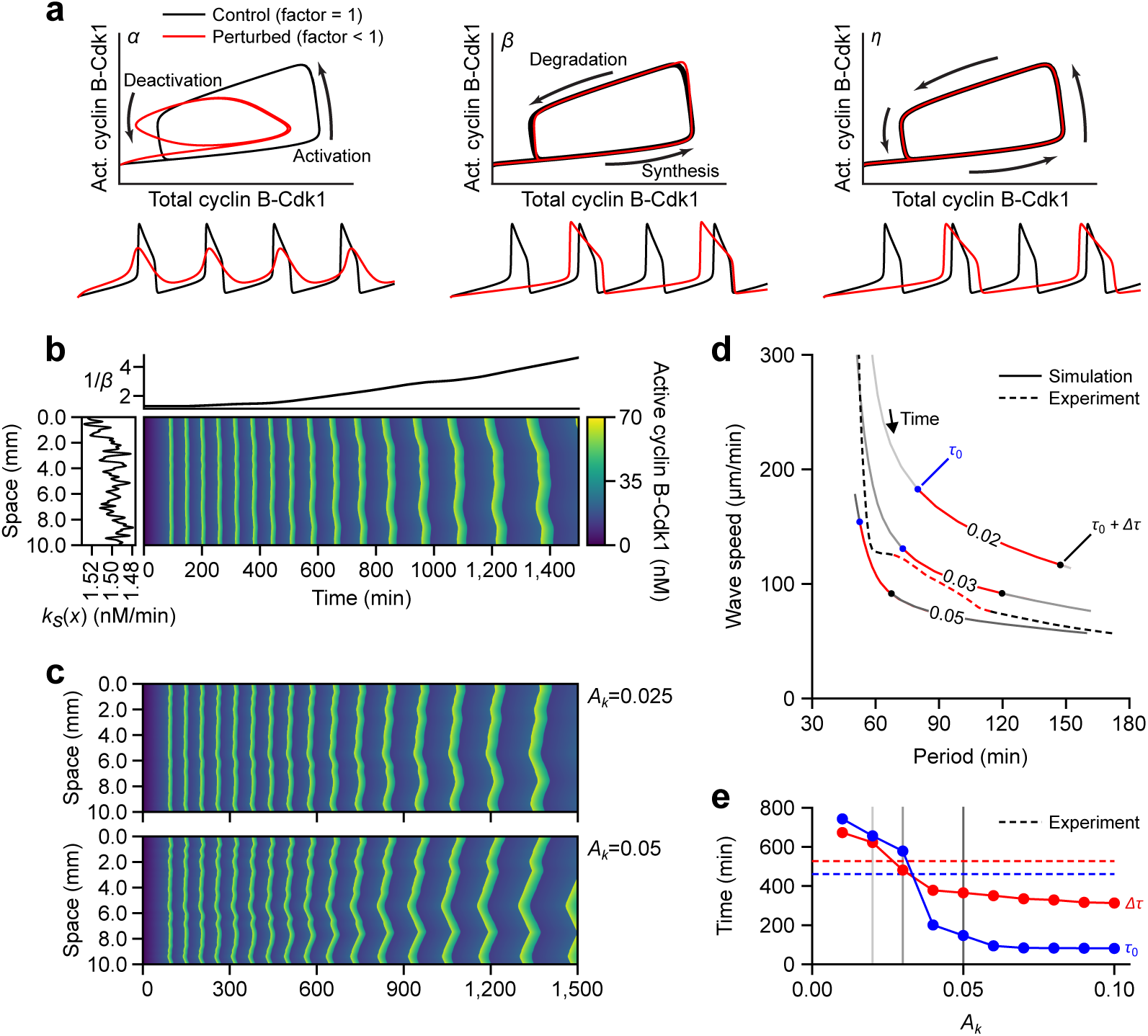
Mathematical modeling explains the transition from phase to trigger waves. **a** Schematic representation of the mitotic dynamics influenced by scaling factors *α* (left), *β* (middle), and *η* (right), which scale for the rates of respective reactions indicated by black arrows. See Methods and Fig. S3 for details. Unperturbed (scaling factor = 1, black lines) and perturbed (scaling factor *<* 1, red lines) dynamics are compared for phase-plane trajectories of total and active cyclin B-Cdk1 concentration (top) and active cyclin B-Cdk1 concentration time courses (bottom). **b** Spatiotemporal evolution of the cyclin B-Cdk1 activity showing the transition from fast to slow waves. Simulation has incorporated the experimental time-dependence of the period shown as 1*/β* (*t*) (top panel) and spatial variability in the synthesis term *k_s_*(*x*) (left panel, see also Methods). **c** Influence of spatial heterogeneity (*A_k_*) on the wave speed entrainment. **d** Speed-period relation of the experiment (dashed line) and the numerical simulations (solid lines with respective *A_k_* values labeled). The transition points *τ*_0_ and *τ*_0_ + Δ*τ* are marked as in Fig. 1e. The time frames between *τ*_0_ and *τ*_0_ + Δ*τ* are highlighted in red. **e** Dependence of transition time scales *τ*_0_ (blue) and Δ*τ* (red) on spatial heterogeneity. Experimental measurements are given in dashed lines. Vertical lines correspond to simulation conditions illustrated in **d** with matching grayscale colors.

To model relaxation dynamics, we incorporate a heterogeneous spatial profile of the cyclin synthesis rate *k_s_*(*x*). The loci with smaller periods in this profile (corresponding to larger values of *k_s_*(*x*)) can act as trigger wave sources, as explored in previous works for pointlike sources^21,22^.

Thus, we perform numerical simulations incorporating these two contributions: a time-dependent *β* (*t*) (Fig. 2**b**, top) and a noisy *k_s_*(*x*) distributed across space (Fig. 2**b**, left). The diffusion rate is set to be 240 µm^2^/min, compatible with a rate reported in the fly embryo^9^, to reproduce the wave speed at the end of the extract lifetime (≳ 1, 000 min). The simulated kymograph in Fig. 2**b** captures both the period lengthening of waves and the gradual formation of slow linear fronts that resemble the qualitative observations from experiments. The formation of linear fronts is influenced by the noise level, denoted by *A_k_* (see Methods for the definition); comparing two simulations with different levels of spatial noise, we find that large heterogeneity (*A_k_* = 0.05) entrains the system more rapidly than small heterogeneity (0.025), leading to a faster reduction to a wave speed that is characteristic of a trigger wave (Fig. 2**c**).

Applying the same analysis used for experimental data allows us to plot the speed-period relation for various heterogeneity levels (Fig. 2**d**). Keeping in mind that the period is a monotonically increasing function of time, we observe that the transition from phase waves to trigger waves happens both earlier (*τ*_0_, blue dots) and faster (Δ*τ*, red regions, end with black dots) as the heterogeneity rises. *A_k_* = 0.03 gives a result that quantitatively agrees with experiments, characterized by a two-step decay with *τ*_0_ = 579 min and Δ*τ* = 482 min.

Finally, we analyze the dependence of the transition time scales on the level of heterogeneity *A_k_* (Fig. 2**e**). Increasing spatial heterogeneity significantly speeds up the transition to trigger waves, occurring both earlier (Fig. 2**e**, blue) and faster (Fig. 2**e**, red). Additionally, we find that suppressing the period elongation does not affect the speedup resulting from increased spatial heterogeneity (Fig. S4). Therefore, the dominant effect in the transition from fast phase waves to slow trigger waves is the finite relaxation time required by traveling waves to establish themselves. All in all, our modeling suggests that this transition can be sped up by introducing spatial heterogeneity into our system.

### Nuclei speed up the transition from phase waves to trigger waves

The question then arises: how can we incorporate spatial heterogeneity in our experimental system? One possible approach is to introduce nuclei into the system. Nuclei can drive wave formation by acting as pacemakers^13,14^. Modeling work demonstrates that the inclusion of pacemakers, whether explicitly or implicitly, drives waves at a frequency that correlates with pacemaker activity, with wave speed being dependent on this driving frequency^8,13,21–23^. Therefore, compartmentalizing the cytosol by introducing nuclei might affect how fast the system transitions from phase to trigger waves.

We supplement extracts with demembranated *Xenopus* sperm DNA (+XS), a method commonly used in the field to reconstitute nuclei^8,13–15^. A representative kymograph of this experiment shows clear wavefronts spanning the whole length of the tube (Fig. 3**a**, top). The zoomed-in region shows individual nuclei forming during interphase, importing active Cdk1 prior to NEB, and disappearing upon NEB (Fig. 3**a**, bottom). Even at this relatively coarse timescale (at 5-min time intervals), we observe the pacemaker nucleus accumulating more active Cdk1 than its neighbors and thereby undergoing NEB earlier (Fig. 3**a**, bottom, red arrows). After NEB, active Cdk1 fills the local region, and pulse-like waves propagate in both directions (Fig. 3**a**, bottom). Similar to the case without nuclei, the spatial profiles clearly indicate that at early times, the patterns resemble phase waves and at later times, trigger waves (Fig. 3**b**; Mov. 2). In other words, despite nuclei visibly forming at early times, we observe a comparable sweeping up of activity, though the effect is much noisier and punctuated by peaks associated with the nuclei themselves throughout the tube (Fig. 3**b**, top). As time progresses, trigger waves become dominant, and the system exhibits clear traveling pulses from the dominating pacemaker nucleus (Fig. 3**b**, bottom).

**Fig. 3.**
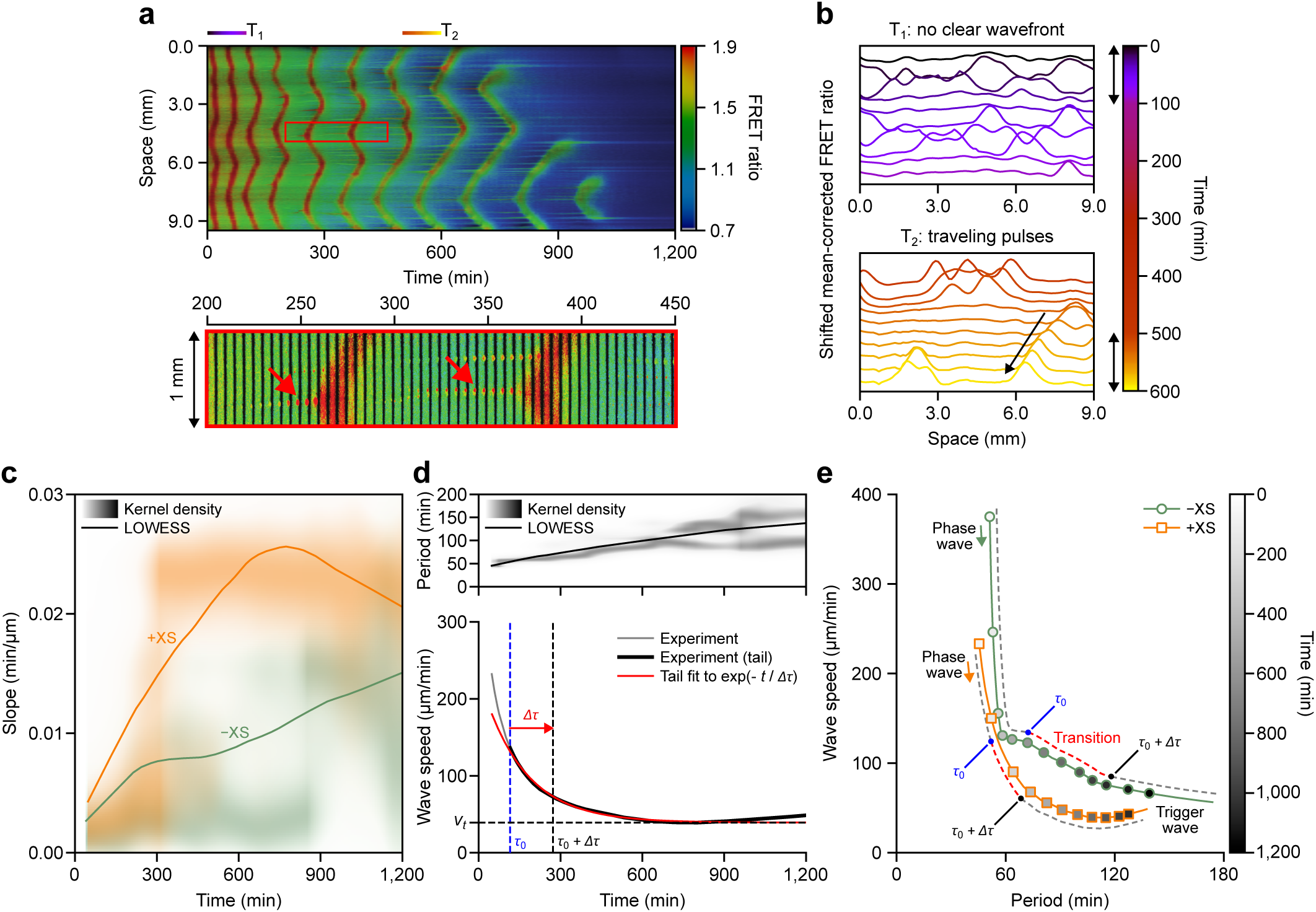
Nuclei entrain the system to the trigger wave regime. **a** Representative kymograph of the effects of adding sperm DNA (+XS). A magnified view of the region inside the red rectangle is provided in the lower panel to show nuclear growth before the entry into mitosis (red arrows). *T*_1_ and *T*_2_ mark the regions used in panel **b**. **b** Comparison of shifted mean-corrected FRET ratio at early times with no clear wavefront (top) and distinctive traveling pulses at late times (bottom). **c** Comparison of the time dependence of the slope for the case of added sperm DNA (+XS, orange) and control (*−*XS, green). Kernel density estimations (normalized at each time) and LOWESS curves are given in the corresponding colors. **d** Time evolution of the period (top) and wave speed (bottom) for +XS. **e** Speed-period relations for both conditions are obtained from LOWESS curves in Fig. 1d (*−*XS) and Fig. 3d (+XS), respectively. Open circles (*−*XS) and open squares (+XS) along the line highlight multiples of 100 minutes. The transition points are marked on dashed guides for each, as done previously.

Repeating the same workflow described previously for this compartmentalized system, we find qualitatively similar behavior for each of the relevant quantities: the slope (Fig. 3**c**) and period (Fig. 3**d**, top) both increase over time, while the maximum activation rate remains constant (Fig. S2**c**). Despite the time evolution of the period being similar between the two conditions (*±*XS), the slope for waves with nuclei consistently exceeds that of waves without nuclei within the 1200-minute observation window (Fig. 3**c**), indicating a potential impact of nuclei on the wave propagation dynamics. To quantify this, we again measure the entrainment time from the exponential fitting (Fig. 3**d**, bottom). We obtain an entrainment time of *τ*_0_ = 115 min and Δ*τ* = 157 min, which is faster than that of the non-nuclei system, indicating a speedup in the transition to trigger waves caused by the presence of nuclei. Interestingly, the fit for both conditions produces a terminal speed close to 40 µm/min, suggesting a common long-term behavior that agrees with speeds previously reported^8,13^. The difference in the entrainment time between the two systems is made clearer when considering the speed-period relation (Fig. 3**e**). As shown, the transient phase waves give way to trigger waves much more rapidly than in the systems without nuclei, as indicated by a decay in speed that begins earlier. The slight increase in speed at late times in the nuclei case is likely due to extract death. Despite this, it is clear that the addition of sperm DNA (nuclei) causes the system to admit trigger waves earlier in time, but also “earlier” in terms of period. This reinforces the idea that wave speed changes due to transient effects, rather than being driven by changes in period.

Our results show that nuclei speed up the entrainment of the system to the trigger wave regime. Nuclei act as pacemakers^13,14^, providing nucleation points for singular wavefronts. Supported by our computational analysis, we extend this notion to argue that the nuclei play a broader role in bringing the system out of the transitory, less-specified phase wave regime and into the well-defined, classical trigger wave regime. In systems without nuclei, patterns remain diffusive and exhibit fast speeds. Over time, trigger waves do develop, albeit slowly. Conversely, systems with nuclei develop trigger waves earlier and more frequently. Consequently, the former displays fast speeds that slowly decrease, while the latter displays speeds that quickly decay and follow a trigger wave speed-period relation.

### Driving waves with metaphase-arrested extract speeds up the transition to trigger waves with and without nuclei

To further understand the entrainment of mitotic waves, we set out to drive waves explicitly by a cytostatic-factor (CSF) extract, similar to a previous study that drove an apoptotic signal through a tube of interphase extract using a reservoir of apoptotic-arrested extract^24^.

CSF extract, a metaphase-arrested extract, is derived from inactivated eggs arrested at meiosis-II^25,26^. While the biological details of CSF arrest remain to be elucidated, the field largely agrees that the Emi family of proteins plays a major role by inhibiting APC/C, with other studies also highlighting the involvement of the Mos-MAPK pathway in CSF arrest^27,28^. Despite these uncertainties, CSF extracts consistently exhibit and maintain high Cdk1 activity unless released from arrest^29^. Moreover, these extracts can be frozen and stored for many months, providing a reliable source of stable high-Cdk1-activity extract^30^. Due to the self-promoting activity of Cdk1 in the mitotic circuit, supplementing oscillating extract with CSF is expected to result in forced activation and consequent propagation of traveling mitotic waves (Fig. 4**a**).

**Fig. 4.**
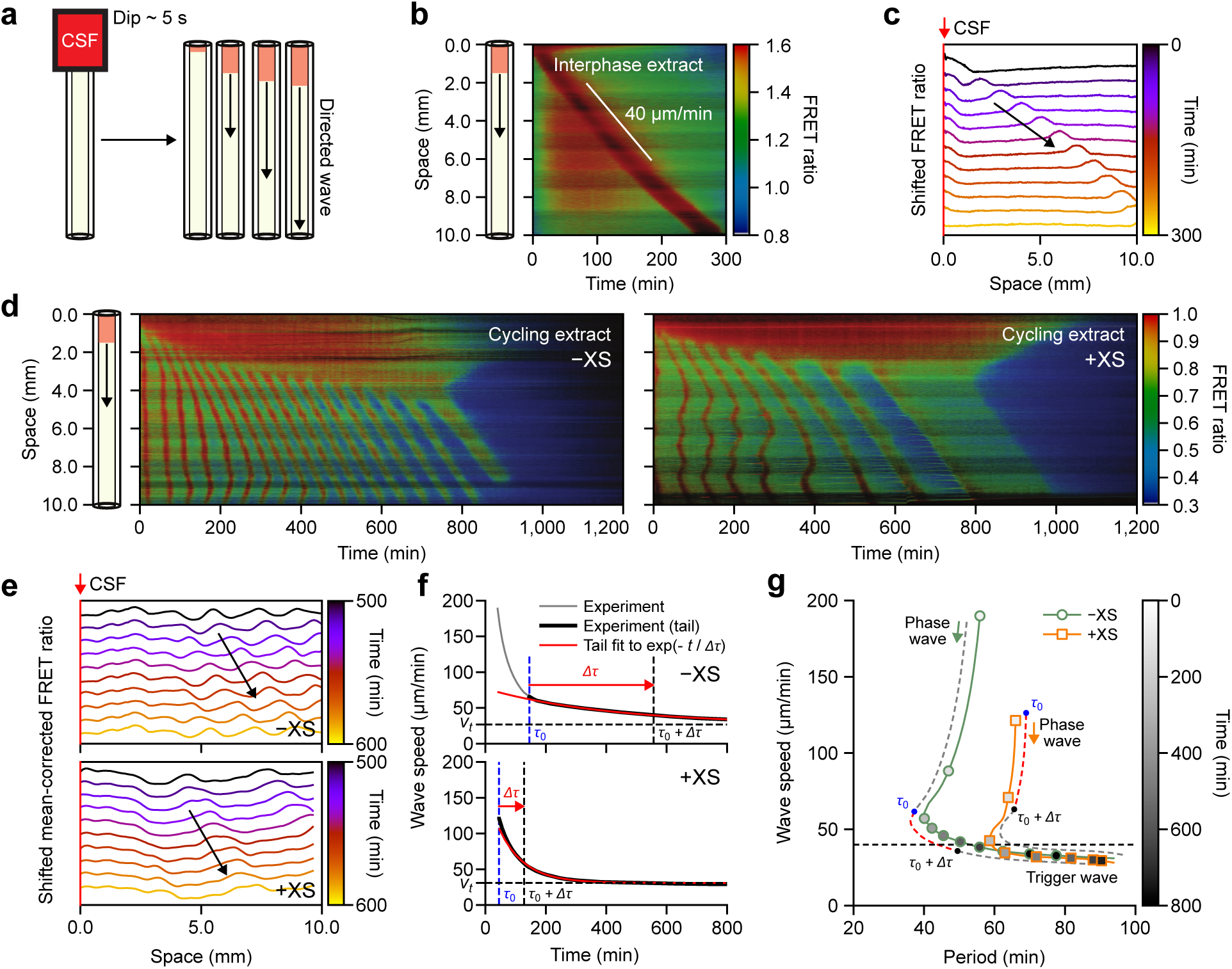
CSF boundary-driven mitotic waves. **a** Schematic representation of the experimental setup to trigger boundary-driven mitotic waves by a 5 second dip in CSF extract. **b** Solitary pulse of high Cdk1 activity propagating with a speed of 40 µm/min (wavefront indicated by a white line) in an interphase-arrested extract triggered by CSF dipping. **c** Spatial profiles of the shifted FRET ratio from the kymograph in **b** showing the excitable pulse. The red vertical line corresponds to CSF-induced arrest. **d** Kymographs for boundary-driven traveling waves in cycling extracts without (left, *−*XS) and with nuclei (right, +XS) present. **e** Traveling waves from kymographs in d revealed by the shifted mean-corrected FRET ratio without (*−*XS) and with nuclei (+XS). **f** Wave speed as a function of time for the two conditions in panel d analyzed via later-time exponential fits (red lines). **g** Speed-period relationship combining both LOWESS estimations for conditions with (+XS) and without (*−*XS) nuclei. Transition points and guides are as previously defined.

To validate this setup, we create a bistable traveling wave using CSF extracts and interphase extracts. Both extracts are prepared following standard protocols in the field^29–31^. We observe that briefly dipping an interphase-extract-filled tube into a CSF reservoir allows the high Cdk1 activity in CSF extracts to excite a traveling pulse of Cdk1 activity in the tube (Fig. 4**b**; Mov. 3). Waves propagate at a consistent speed of 40 *±* 2 µm/min, in agreement to previously measured mitotic trigger waves in cycling extracts^8,13,14^. Visualizing the spatial profiles by shifting the FRET ratio reveals a clear traveling peak of activity, confirming the presence of trigger waves (Fig. 4**c**). This experiment demonstrates the efficacy of using CSF extracts as an explicit source to drive waves.

We thus applied the CSF extracts to drive tubes filled with cycling extract with or without sperm DNA added; in both cases, we observed the mitotic arrested region persistently drives multiple cycles of Cdk1 activity waves throughout the tube (Fig. 4**d**; Mov. 4 and Mov. 5). Similar to the non-driven experiments, wavefronts appear as pulses of high Cdk1 activity, regardless of the presence of nuclei, validating the emergence of trigger waves (Fig. 4**e**).

Applying the same analysis pipeline as before, we study how period and wave speed change over time. Initially, both conditions show a slight decrease in the oscillation period, likely due to the diffusing influence of CSF, followed by a “typical” period elongation (Fig. S5**a**). While the observed behavior is qualitatively similar, the two conditions show a difference in the magnitude of the change in period despite experiencing the same driving force (Fig. S5**b**). When measuring the wave speed in terms of slopes, we find that the system with nuclei has higher values than their non-nuclei counterparts (Fig. S5**c**), matching what we observe in the non-driven system (Fig. 3**c**). This is likely due to the longer periods in the system with nuclei, but also suggests a possible difference at the level of wave propagation.

Our time scale analysis of wave speed reveals that the entrainment time for the condition with nuclei (+XS, *τ*_0_ = 45 min, Δ*τ* = 83 min) is shorter than without (*−*XS, *τ*_0_ = 146 min, Δ*τ* = 411 min) (Fig. 4**f**). The fact that entrainment times for driven waves are significantly shorter than their non-driven counterparts (Fig. 4**f**, *±*XS, as compared to Fig. 3**d**, *±*XS) seems to point to a cumulative effect from multiple pacemakers: both the CSF and nuclei contribute to entrainment. Interestingly, the entrainment time for the CSF-driven case without nuclei (*−*XS/+CSF, *τ*_0_ = 146 min, Δ*τ* = 411 min) is still longer than the undriven case with nuclei (+XS*/ −* CSF, *τ*_0_ = 115 min, Δ*τ* = 157 min). It is plausible a set of multiple, but theoretically weaker, pacemakers could entrain the system faster given a distributed effect throughout space. In addition, the relaxation time scale for the CSF-driven case without nuclei (Δ*τ* = 411 min) is not shortened significantly compared to the non-driven experiment (Δ*τ* = 528 min), although CSF driving brings systems to initiate the transition more quickly (*τ*_0_ = 146 and 462 min, respectively). In contrast, the presence of nuclei shortens both the initiation time and the relaxation time. This discrepancy underscores a fundamental biological difference in how CSF and nuclei contribute to entrainment, which is worth further investigation. Additionally, the fitted terminal speed 30 µm/min matches what we and others observed previously^8,13,14^.

Importantly, driving the system in this way explicitly entrains the system to the trigger wave regime, quickly and permanently. The phase waves of early times start to transition to trigger waves within two or three cycles and propagate across the entirety of the tube. These waves appear to follow a clear speed-period relation, distinct from the undriven case (Fig. 4**g**). In both cases, we see a fast decrease in speed with small changes in period that lead to a smooth approach to terminal speed (Fig. 4**g**). In this way, driving the system elucidates a clear difference between the transients—and possible phase waves—of early times and trigger waves, and reinforces the notion of entrainment explicitly.

### The presence of phase waves and trigger waves depends on spatial heterogeneities in initial conditions and system parameters

There is a key conceptual distinction between phase waves and trigger waves. While information transmission by phase waves is nearly absent and synchrony is only maintained when the initial phase difference is small, trigger waves transmit information over long distances. The mechanisms underlying the two kinds of waves are also different. Although both appear in oscillatory systems, trigger waves require timescale separation and spatial coupling commonly mediated by diffusion. Timescale separation in our experiments is naturally present due to the rapid activation of cyclin B-Cdk1 compared to cyclin buildup from synthesis. It is this rapid activation that excites neighboring regions through protein diffusion and triggers a sustained propagation of the wave. In contrast, phase waves appear as a result of a small delay in the activation time of adjacent positions, creating a structured phase difference that takes the appearance of a wave. Another important difference is in the speed of the wave and its stability. The speed of a trigger wave is uniquely defined by the properties of the underlying oscillator and diffusion, and it is said to be stable because waves with different propagation speeds will converge to the stable one. Conversely, a phase wave is not stable, it can appear at any speed, and diffusion will attenuate phase differences with time until the system oscillates synchronously.

To demonstrate the difference, we use numerical simulations to initiate a trigger wave by introducing a period difference at the center of the spatial domain with spatially homogeneous levels of activity as the initial condition (Fig. 5**a**, (i)). In contrast, we initiate a phase wave by keeping the period fixed in space and asserting an initial condition where there is a linear difference in the phase of the oscillator that decreases as one moves far from the central position (Fig. 5**a**, (ii)). Then we explore the robustness of both systems by reducing the timescale separation of the oscillator (Fig. 5**a**, (iii, iv)), where the timescale separation is controlled by factor *α* for relaxation-like (*α* = 1, Fig. 2**a**, left, black lines) and sinusoidal (*α <* 1, Fig. 2**a**, left, red lines) oscillations. Quantifying the shapes of wavefronts at late times with the fit *x* = *d* + *vt^γ^*, we obtain the dependence of the trigger-wave-likeness on *α*, summarized in Fig. 5**b**. We found that only when the oscillations are relaxation-like, indicated by larger values of *α*, trigger waves establish in space in a stable manner, entraining the oscillatory background and displaying a linear front (*γ* = 1, Fig. 5**a**, (i)), while low values of *α*, which correspond to sinusoidal oscillations, lead to a curved front (*γ >* 1, Fig. 5**a**, (iii)). Increasing timescale separation with *α* also amplifies the penetration depth of the trigger wave into the medium^21,22^, as shown in the long-time behavior, comparing Fig. 5**a**, (i) and (iii). On the other hand, phase waves do not change with the timescale separation, comparing Fig. 5**a**, (ii) and (iv). The spatially averaged wave speed, measured at the wavefront segments in the kymograph (Fig. 5**a**, (i), red lines), converges to the stable value expected for trigger waves (Fig. 5**c**, (i)); in contrast, the speed for phase waves (Fig. 5**a**, (ii), red lines) increases with time as the oscillations synchronize (Fig. 5**c**, (ii)).

**Fig. 5.**
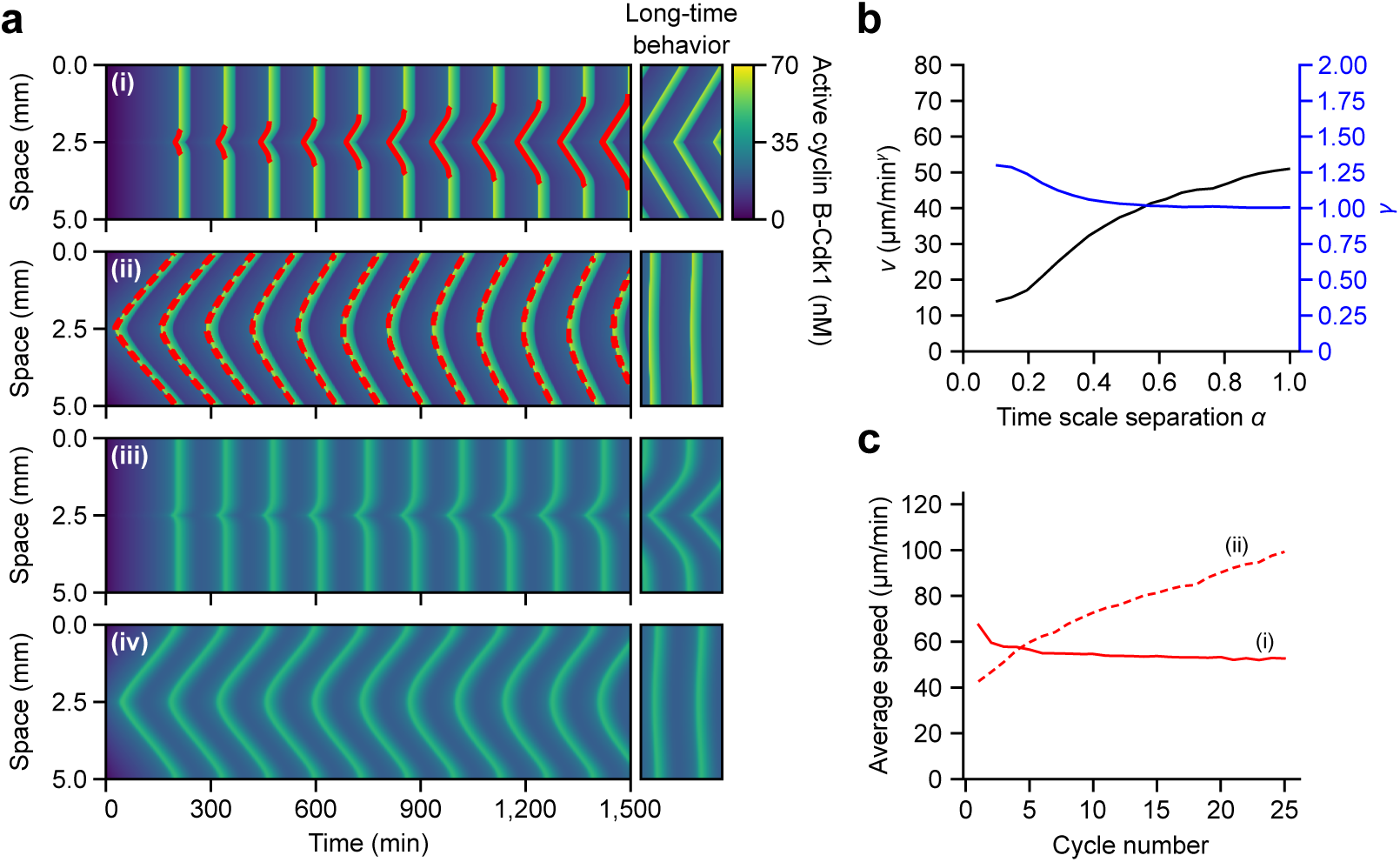
Influence of the oscillator properties and spatial heterogeneity on the formation of phase and trigger waves. **a** Kymographs representing the spatiotemporal evolution of cyclin B-Cdk1 activity. (i) Simulation with a spatially homogeneous initial condition of activity and a spatially heterogeneous period dependence to introduce a pacemaker at *x* = 2.5 mm, exhibiting trigger waves for *α* = 1. (ii) Simulation with a spatially linear phase difference in the initial condition of activity and a spatially homogeneous period, exhibiting phase waves for *α* = 1. (iii) Same spatial heterogeneity as (i) for *α* = 0.1. (iv) Same spatial heterogeneity as (ii) for *α* = 0.1. **b** *v* and *γ* as functions of *α* resulting from fitting the long-term shapes of pacemaker-driven waves in **a** with the expression *x* = *d* + *vt^γ^*, showing a progressive transition to linearly propagating trigger waves (*γ* = 1) with a stable speed. **c** Temporal evolution with cycle number of the spatially averaged wave speed for phase and trigger waves in the shown kymographs (i) and (ii), panel **a**. Speed is measured only at the wavefront segments indicated in red lines.

Together, our numerical simulations (Figs. 2 and 5) suggest a crucial role of spatial heterogeneity in the phase-to-trigger wave transition, confirmed by our experimental observations, which we recapitulate in Fig. 6. First, disrupting oscillator homogeneity in space makes the system lose synchrony earlier in time (Fig. 6**a**). This acceleration is quantified in terms of the transition starting time *τ*_0_. *τ*_0_ is greatly reduced by introducing multiple nuclei in the system (115 min, +XS), driving the system with CSF extract from the boundary (146 min, +CSF), or combining both of them (45 min, +XS/+CSF), compared to the control experiment (462 min, Control). The transition rate is also affected substantially by heterogeneity, particularly by nuclei (Fig. 6**b**). The slowdown of wave speed after *τ*_0_ that approaches the terminal trigger wave speed is well characterized by an exponential decay with a time scale Δ*τ* that varies across experimental conditions. Δ*τ* is much shorter for extracts with nuclei (157 min for +XS and 83 min for +XS/+CSF) as opposed to cytoplasm-only experiments (411 min for +CSF and 528 min for Control), which signifies the important role of having multiple pacemakers in the entrainment.

**Fig. 6.**
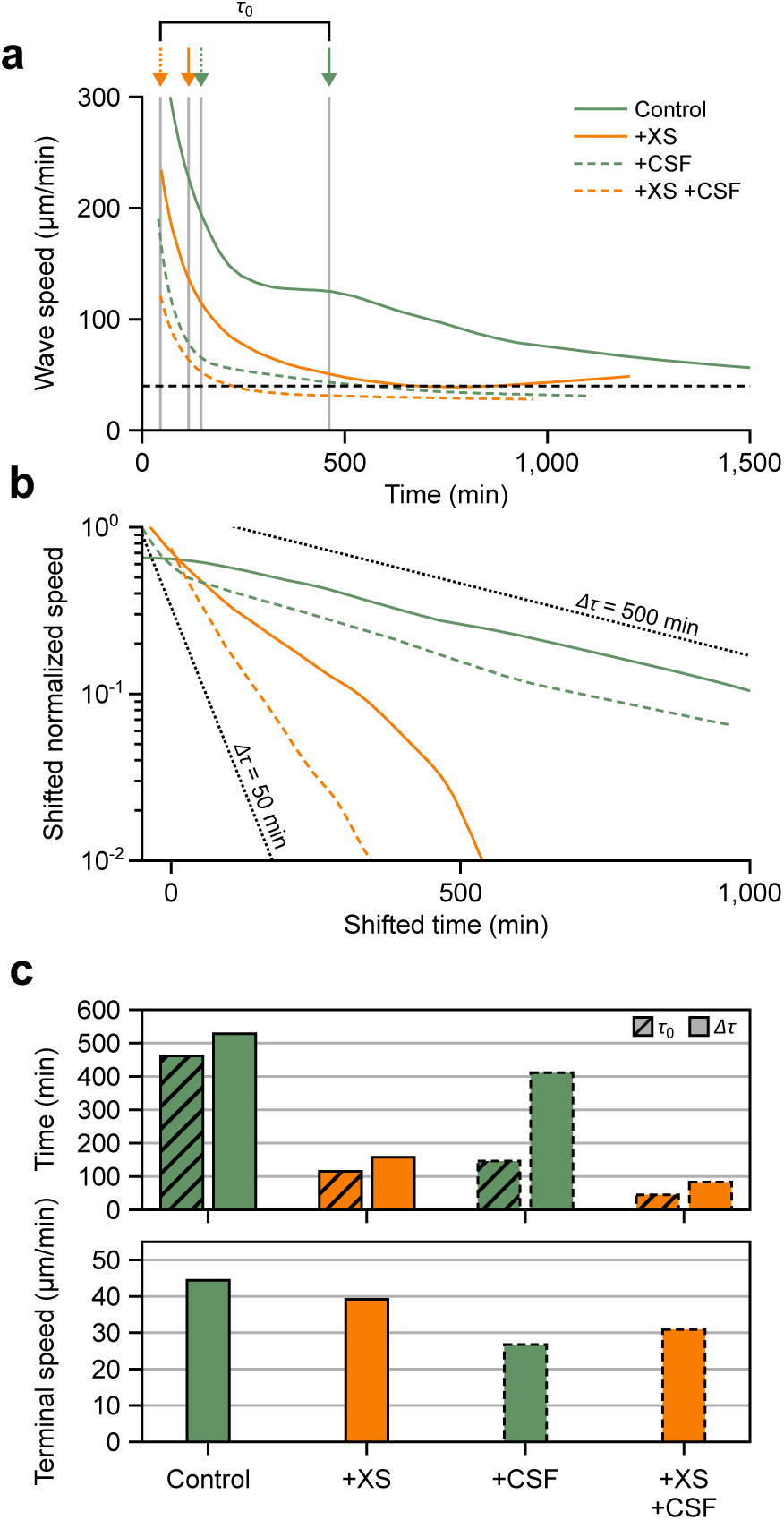
Spatial heterogeneity coordinates phase-to-trigger wave transition. **a** Time evolution of wave speed. Each vertical line with the corresponding arrow on the top indicates *τ*_0_ for each experimental condition (462, 115, 146, and 45 min for Control, +XS, +CSF, and +XS/+CSF, respectively). The horizontal line depicts the terminal for the excitable system (40 µm/min). **b** Exponential relaxation of late-time wave speed. Time is measured from respective *τ*_0_, and speed is offset by the terminal speed and then normalized against the speed at *τ*_0_. The negative reciprocal of the curve slope gives a visual estimation of Δ*τ*, which are 528, 157, 411, and 83 min for each condition. Dotted guidelines correspond to 50 and 500-min time scale relaxations, respectively. **c** Transition time scales *τ*_0_ and Δ*τ* (top) and terminal speeds (bottom). Terminal speeds are 44, 39, 27, and 31 µm/min for each condition, converging to a similar level and comparable to the traveling speed of the activation pulse in interphase extracts driven by CSF (40 µm/min).

Interestingly, despite the spatial heterogeneity may cause one-order-of-magnitude changes both in *τ*_0_ and Δ*τ*, the terminal speed across all conditions remains mostly unchanged (Fig. 6**c**). This suggests that regardless of the specific mechanism that drives the transition, coupled mitotic oscillators eventually synchronize with a consistent timing gradient established by trigger waves of 30-40 µm/min speed. A recent work by Huang *et al*. also highlighted the robustness of mitotic trigger wave speed under the physical stress of changing cytoplasmic concentrations^32^. Such a particular feature of being a reliable reference of timing distinguishes trigger waves from phase waves. In the absence of dynamic constraints imposed by diffusion, phase waves propagate at a fast but arbitrary speed that may depend on different physiological circumstances individual cells face. The speed of trigger waves, on the other hand, is a more intrinsic property of a dynamical system as explored in our theoretical work (Fig. 5).

## DISCUSSION

Spatial coordination is essential to communicating complex biological processes. In this work, we probed the nature of one such coordination mechanism: mitotic waves that coordinate the process of cell division in large cells. Using a frog egg extract system which reproduces cell cycles *in vitro*, we characterized how mitosis spreads through the *Xenopus laevis* cytoplasm via either phase waves or trigger waves.

Although properties of trigger waves have been thoroughly studied in the literature^21,22,33–37^, the properties of their transient dynamics have not. Through our frog egg extract experiments in thin long (quasi-one-dimensional) tubes, we observed phase waves in the transient dynamics towards the formation of a stable trigger wave. Even though cell cycle oscillations also slowed down, we showed that this is not required to observe a transition from phase waves to trigger waves. Certainly, it can take a long time before a pacemaker—a region that oscillates faster than its surroundings—is able to entrain its surroundings via trigger waves^21,22^. While in excitable media, a trigger wave can travel uninterrupted throughout the medium, in oscillatory media the entrainment distance is limited by the inherent oscillatory period of its surroundings. Even in the transient time when a trigger wave is still forming, the regular biochemical oscillations that drive the early embryonic cell cycle in *Xenopus laevis* will drive the whole system into mitosis throughout the whole medium. Any phase gradients will thus give rise to phase waves as the cell cycle phase is swept up.

This puts our work in dialogue with the existing literature regarding mitotic waves in *Drosophila*. Although period lengthening was found to be a driver of a sweep-to-trigger transition in *Drosophila*^17^, we showed that this is not required for directing a transition from phase waves to trigger waves. Furthermore, this and other work highlight that phase wave properties are not robust to heterogeneity as they do not correspond to an actual attracting system solution. In Hayden *et al*.^17^, the authors also show embryos displaying a nuclear density gradient in the syncytial embryo can lead to trigger rather than sweep or phase waves. We demonstrate a similar effect of disrupting homogeneity both by adding nuclei to the homogeneous system and by driving it explicitly with CSF. In all three cases, these trigger-wave-producing effects overtake the underlying quasi-synchronous patterns.

As phase waves do not actively propagate through a medium and require structured initial phase differences over a certain distance, they typically only persist for hundreds of micrometers. A trigger wave can travel long distances (*∼* 10 mm), and the typical length scale of the concentration gradient at the leading front ranges around hundreds of microns as well. This underscores the prevalence of phase waves in relatively small embryos, such as *Drosophila*, due to the limited length to accommodate a trigger wave front gradient. On the scale of some biological functions, the propagation distance of phase waves is relevant. However, for the specific purpose of coordination, this proves insufficient in larger cells. Trigger waves, as made evident by our work here, conversely, transmit signals orders of magnitude farther in distance. This questions the physiological relevance of phase waves. At most, one could argue for a tradeoff between speed and distance. For mitosis it could be reasonable that nature would select for trigger waves in larger embryos such as *Xenopus*, where coordinating over large distances is more relevant than in *Drosophila*. Recently reported ultrafast waves^10,16,38,39^, faster-than-trigger-wave signaling achieved without requiring bistable reactions or diffusion-mediated coupling, necessitate further comprehensive studies to understand the fundamental nature of these waves in comparison to classical trigger waves. Our work highlights the importance of examining not only stable waves but also the time evolution of waves as they develop. It also provides a framework that integrates experiments and theory to dissect the transition between different wave regimes.

Future experiments could expand on this work by pursuing other forms of perturbations by inhibiting the feedback loops in the network. The field already demonstrated the importance of Wee1 for “forming the trigger”^8^, but with our setup and analysis framework, one could quantify the effect and provide stronger evidence in either direction. The same applies to Cdc25, a phosphatase acting antagonistically to the Wee1 kinase in the regulation of Cdk1, both forming positive feedback loops with Cdk1. It would be interesting to observe whether Wee1 and Cdc25 affect this time dependence in similar manners. Moreover, we know that these inputs also translocate in and out of the nucleus throughout one cycle, at different times^40,41^. It stands to reason that inhibition thereof could change in the presence of nuclei, and thus, we might see a nuclei-dependent effect on how inhibition perturbs this transition. Clearly, much work remains on elucidating the details of time-dependent wave behavior.

In another vein, the CSF driving setup could be used to expand on this study by asking how perturbations to the clock network, including inhibitions for other clock constituents such as Cdc25, Wee1, PP2A, etc., change trigger wave propagation. Furthermore, like CSF extracts, interphase extracts maintain activity for months while frozen, making them more accessible for faster and simpler data acquisition than involving the cycling system. In practice, such experiments could provide a more straightforward method for testing all of the perturbations mentioned above: clock inhibitors, glycerol-modulated diffusion, etc. In particular, this would facilitate a direct examination of whether nuclei indeed perturb wave propagation as it would eliminate their dual role as a pacemaker. In total, this setup offers a wealth of opportunities to probe trigger wave dynamics, relevant for *in vivo* embryogenesis in *Xenopus*.

Moreover, one can envision perturbing the source itself. Theory predicts the wave speed to depend on the difference between the pacemaker and bulk frequency^21,22^. Modulating the driving force of the CSF source, whether through dilution or the use of inhibitors, can provide a direct test of these theoretical predictions. As the interplay between CSF arrest and its driving force remains unclear, comprehending such perturbations requires additional modeling efforts. Nevertheless, our successful demonstration of driving waves *in vitro* underscores the significance of elucidating these interactions. Taken together, these future investigations would not only enhance our understanding of how organisms transmit mitotic information across long distances, but also provide fundamental insights into the nature of biochemical waves generally, and phase waves and trigger waves in particular.

## METHODS

### Xenopus laevis egg extracts

To capture mitotic waves *in vitro*, we made cell-free cycling extracts from *Xenopus laevis* eggs following a published protocol^18,19^ adapted from Murray^26^. Extracts were then supplemented with various reporters, drugs and/or sperm DNA, depending on the experimental conditions. The Cdk1-FRET sensor was prepared as described in Maryu and Yang^15^. Demembranated sperm DNA was prepared following the established protocol^26^. Work from the Yang lab demonstrated an intermediate range of dilution of the extracts can improve the number of cycles, with the best activity at around 20% dilution^42^. As a result, for the data described here, the dilution was kept constant at 20% with extract buffer (100 mM KCl, 0.1 mM CaCl_2_, 1 mM MgCl_2_, 10 mM potassium HEPES, 50 nM sucrose, pH 7.8). Extracts were subsequently loaded into 5-10 mm-long sections of Teflon-coated Masterflex PTFE tubing (inner diameter 150 µm) via aspiration, submerged under mineral oil, and then recorded using time-lapse epifluorescent microscopy (Olympus IX-83). Given the dimensions of the tubing used, tubes were organized into, at most, groups of five in one direction. We did not observe any significant contamination between tubes, even when forcing them into such close proximity.

For Figs. 4**b** and 4**c**, frozen interphase extracts were made using the standard protocol in the field^20,31^. On the day of the experiment, one aliquot is thawed on ice, supplemented with reporters, and then loaded into PTFE tubing for imaging.

The CSF extracts were made following established protocols^29^ adapted from the original^26^. Using laid eggs, we produced large quantities of extract (on the order of mL), vastly more than necessary for a single experiment (10-20 µL). To preserve said large quantities of extract, we implemented a freezing protocol adopted from Takagi and Shimamoto^30^. To maintain conditions across the reservoir and the cycling extract, CSF extracts were also diluted to 20% with extract buffer. No reporters or drugs were added to these extracts.

In order to set up the CSF driven system, we first cut PTFE tubing into individual sections of *∼* 10 mm lengths and loaded each via aspiration such that the extract (either interphase or cycling extracts) filled the tube in excess: visual inspection of the syringe adapter showed the fluid line exceeding the tube opening. Then the tube was dipped into the CSF reservoir syringe-end first for 5-10 seconds to ensure fluidic contact between the cycling/interphase and CSF extracts. While the original apoptotic wave paper^24^ described maintaining contact between the reservoir and tubes for many minutes, we observed any contact longer than *∼* 10 seconds resulted in mitotic arrest overtaking most, if not all, of the tube. This sometimes occurred even at shorter dipping times. As such, care was taken to minimize the contact time. Tubes were then submerged under mineral oil and imaged as discussed above.

### Image processing and analysis methods

Grids of images were captured and subsequently stitched together using ImageJ’s Grid/Pairwise Stitching plug-in^43^, in conjunction with additional pipeline code written in Fiji/Java. Bright-field images from the first frame were used to generate stitching parameters, which were fed to ImageJ to stitch each channel at each frame consecutively. While capturing grids of images in this way resulted in a non-zero time lag between subsequent sections along a tube, and multiple of this lag between the first and last sections, this gap amounted to a few seconds, much smaller than the scale of the overall imaging timestep which was on the order of minutes. As such, this was ignored for the purposes of analysis. The stitched stacks were then straightened using Fiji and a manually selected curve from the bright-field images as an input. This curve was unique to each tube, though the profiles of the tubing sections often followed roughly the same shape, with not much distortion. Afterwards, the tube images were cropped so as to only include the inner dimension, again using the bright-field images as a guide. Additionally, the FRET ratio was calculated separately as in Maryu and Yang^15^.

For the analysis of wavefronts, first, individual kymographs were corrected for any decaying baseline trend, and any NaN pixels were filled using the scikit-image function inpaint. Afterwards, we detected peaks for each time series at each pixel along the tube. The peaks themselves were then clustered into individual cycles in Python. Once cycles were identified and separated, the collection(s) of peaks were fitted and/or smoothed in space and time, after which slopes (and speeds) were calculated along each front by taking the numerical derivative of the fits at each point. Periods followed directly from the detected peaks. The normalized density estimation for the time dependence of the period, slope, and maximum activation rate made use of the SciPy function scipy.stats.gaussian kde using a Gaussian kernel of *σ_t_* = 50 min as a sliding window for time and the maximum value normalized to one.

### Moving horizon fitting

To determine *τ*_0_, we examined the quality of the exponential fitting of the wave speed’s later-time decay, by calculating mean square residual (MSR) of the fitting for the time frame [*τ*_0_, ∞). Both the MSR (Fig. S1, top) and its derivative with respect to *τ* (Fig. S1, bottom) sharply changed (Control, +XS, and +CSF) at a finite *τ* indicating the fitting at the tail segment abruptly worsened upon extending it to earlier times. *τ*_0_ was defined as the largest time that the derivative lies below a chosen threshold, *−*0.025 µm^2^*/*min^3^ (horizontal red line). *τ*_0_ was affected minimally by the choice of the threshold due to the sharp change in the derivative. This definition applied consistently across data with varying fitting quality (for example, +XS data has an overall lower fitting quality than +XS/+CSF data), making it preferable over other thresholding methods based solely on MSR. If the fitting quality is good for all scanned *τ* values (in the case of +XS/+CSF), the earliest recorded time was defined as *τ*_0_.

### Mathematical model

We use the mathematical model previously introduced^8,20^ describing the dynamics of the total cyclin B-Cdk1 *c ≡ c*(*x, t*) and its active form *a ≡ a*(*x, t*). The equations can be written in the following form:

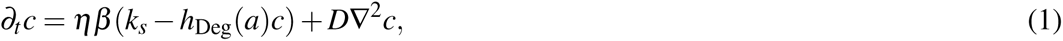

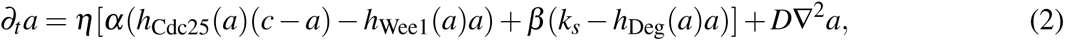

where the newly introduced dimensionless parameters are *α*, *β*, and *η*. The parameter *η* scales all parameter rates and allows for precise control of the period of the oscillations. The parameter *α* scales the rates related to activation-inactivation processes mediated by Cdc25 phosphatase and Wee1 kinase, which are described by Hill functions of the form

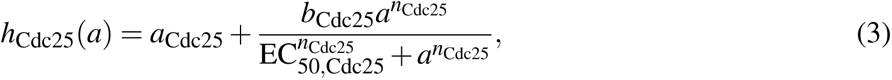

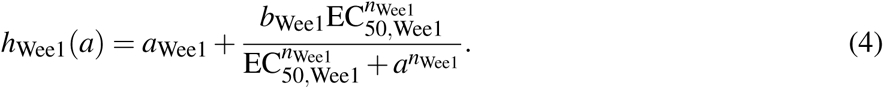

The parameter *β* scales synthesis and degradation rates to control the time spent in interphase and mitosis. The degradation term encompasses the APC/C-induced degradation of cyclin B which is described with the hill function

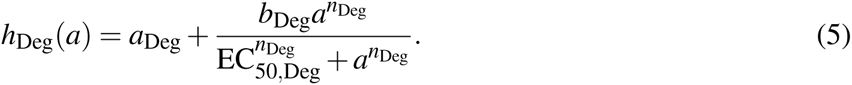

The parameters of the model are *k_s_* = 1.5 nM*/*min, *a*_Cdc25_ = 0.8 min*^−^*^1^, *b*_Cdc25_ = 4 min*^−^*^1^, EC_50_,_Cdc25_ = 35 nM, *n*_Cdc25_ = 11, *a*_Wee1_ = 0.4 min*^−^*^1^, *b*_Wee1_ = 2 min*^−^*^1^, EC_50_,_Wee1_ = 30 nM, *n*_Wee1_ = 3.5, *a*_Deg_ = 0.01 min*^−^*^1^, *b*_Deg_ = 0.06 min*^−^*^1^, EC_50_,_Deg_ = 32 nM, *n*_Deg_ = 17, *α* = *β* = *η* = 1 and are kept constant in this work otherwise specified.

### Numerical simulations

The model described by Eqs. (1) and (2) is a system of two coupled partial differential equations integrated in time with a pseudo-spectral method^44^. We consider a grid with *N_x_* grid points to describe a spatial domain of length *L_x_*with no-flux boundary conditions and we integrate with a timestep Δ*t* the linear terms in Fourier space exactly, while the nonlinear terms are integrated using a second-order in time approximation.

Numerical simulations showing the transition from phase to trigger waves in time (Figs. 2**b**, 2**c**, 2**d**, and 2**e**) have been performed using the integration parameters *N_x_* = 4096, *L_x_* = 10 mm, and Δ*t* = 0.002 min starting with an initial condition of *a*(*x*) = *c*(*x*) = 0 nM introducing spatial heterogeneity in the synthesis term of the following form: *k_s_*(*x*) = *k_s_*[1 + Θ(*x*) + *A_k_N*(*x*)], where Θ(*x*) is a manually introduced profile to induce pacemakers at chosen locations for visualization porpoises and set to zero in Fig. 2**c** to explore the impact of the synthesis noise amplitude *A_k_*. The introduced heterogeneity *N*(*x*) is computed by generating colored noise *n*(*x*) = *F^−^*^1^[exp(*−*(*σk*)^2^*/*2 *−* 2*iπu_k_*)](*x*) using the inverse Fourier transform *F^−^*^1^ where *u_k_* is a random number uniformly distributed between 0 and 1 for each Fourier mode *k* and the typical length scale of the spatial heterogeneities is chosen *σ* = 77.46 µm. The noise is later normalized to have a maximum value of one with the expression,

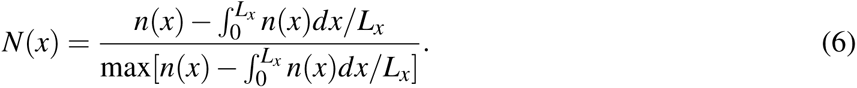

The temporal dependence *β* (*t*) is computed using the linear dependence of the period *T* (1*/β*) shown in Fig. 2**a** (middle) to reproduce the experimentally observed time-dependence of the period in Fig. 1**d** (top). The calculated *β* (*t*) is initialized at the time of the second oscillation, which corresponds to the first measurement of the period in the experiments.

Numerical simulations in Fig. 5 have been performed using the integration parameters *N_x_* = 1024, *L_x_* = 5 mm, and Δ*t* = 0.002 min starting with an initial condition *a*(*x*) = *c*(*x*) = 0 and introducing a pacemaker at the center using a step function as Θ(*x*) = ΔΘ*H*(*s/*2 *−|x −L_x_/*2*|*) where ΔΘ = 0.3, *H*(*x*) is the Heaviside function, and *s* = 50 µm for simulations showing trigger waves. Simulations showing phase waves use a constant value of Θ(*x*) and as initial condition for *a*(*x*) = 2(*a*_min_ *−a*_max_)*|x−L_x_/*2*|/L_x_* + *a*_min_ a linear triangular spatial profile of decreasing activity from *a*_max_ = 20 to *a*_min_ = 0 nM as the distance from the center increases.

## CODE AVAILABILITY

Python codes for the analysis of mitotic waves properties and Fortran codes for performing numerical simulations are deposited on GitHub. Codes are available from Zenodo: https://zenodo.org/doi/10.5281/zenodo.10583185^45^ and from Gelens Lab GitLab https://gitlab.kuleuven.be/gelenslab/publications/mitotic waves.

## ACKNOWLEDGEMENTS

Q.Y. acknowledges funding from the National Science Foundation (MCB#2218083) and the National Institutes of Health (R01GM144584). L.G. acknowledges funding from the Research Foundation Flanders (FWO, grant number G074321N). D.R.-R. is supported by the Ministry of Universities through the “Pla de Recuperació, Transformació i Resilència” and by the EU (NextGenerationEU), together with the Universitat de les Illes Balears.

## SUPPLEMENTARY FIGURES

**Fig. S1.**
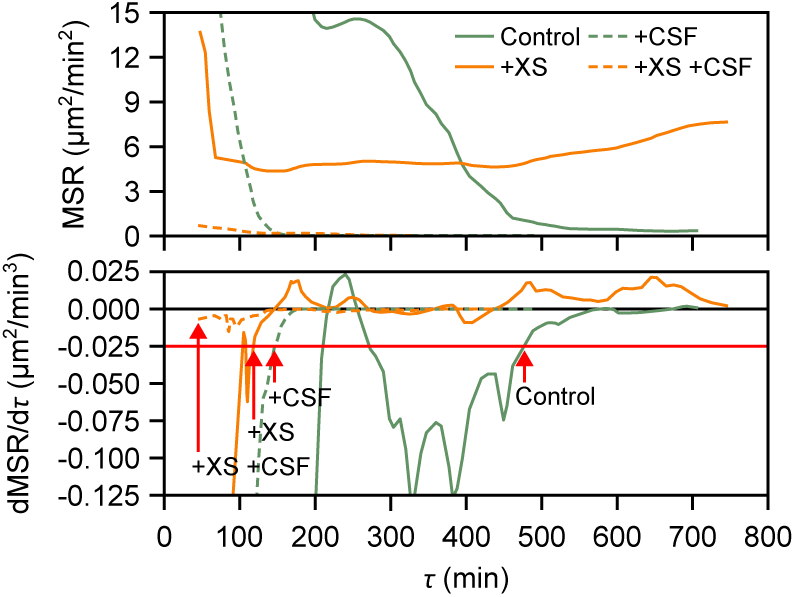
Mean square residual (MSR) of the exponential fitting of the wave speed at [*τ,* ∞) (top), and its *τ*-derivative (bottom). *τ*_0_ is defined as the largest *τ* when the derivative is below a threshold (horizontal red line). *τ*_0_ for each experimental condition is indicated with a red arrow.

**Fig. S2.**
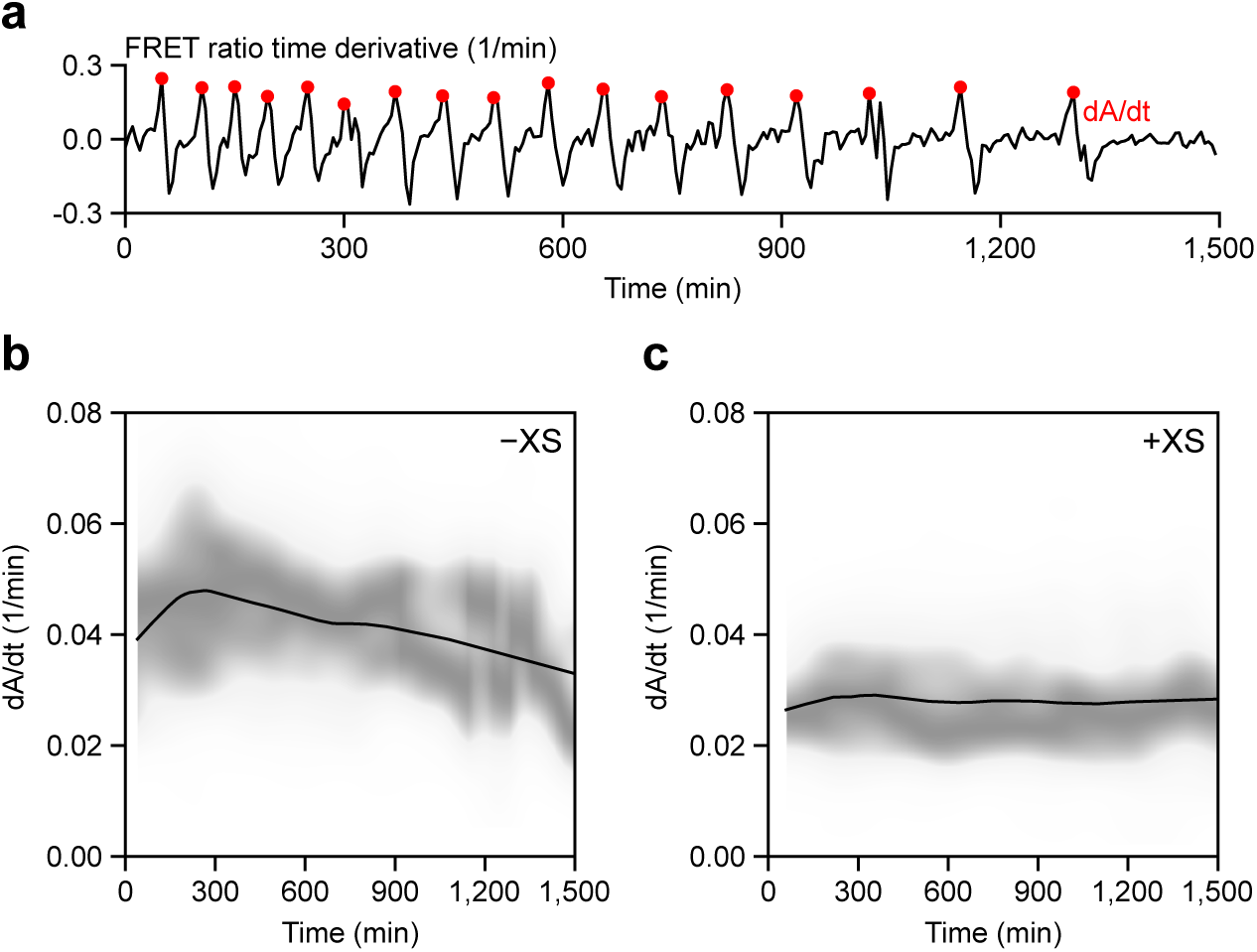
**a** Definition of the maximum activation rate, *dA/dt*. The largest time derivative of FRET ratio per cycle, indicated in red dots, is defined as *dA/dt*. The time course is taken at *x* = 10 mm of the kymograph given in Fig. 1b. **b** Maximum activation rate in non-driven experiments without sperm DNA (*−*XS), represented by the kernel density (gray colormap) and LOWESS (solid black line) estimations. **c** Maximum activation rate in non-driven experiments with sperm DNA (+XS).

**Fig. S3.**
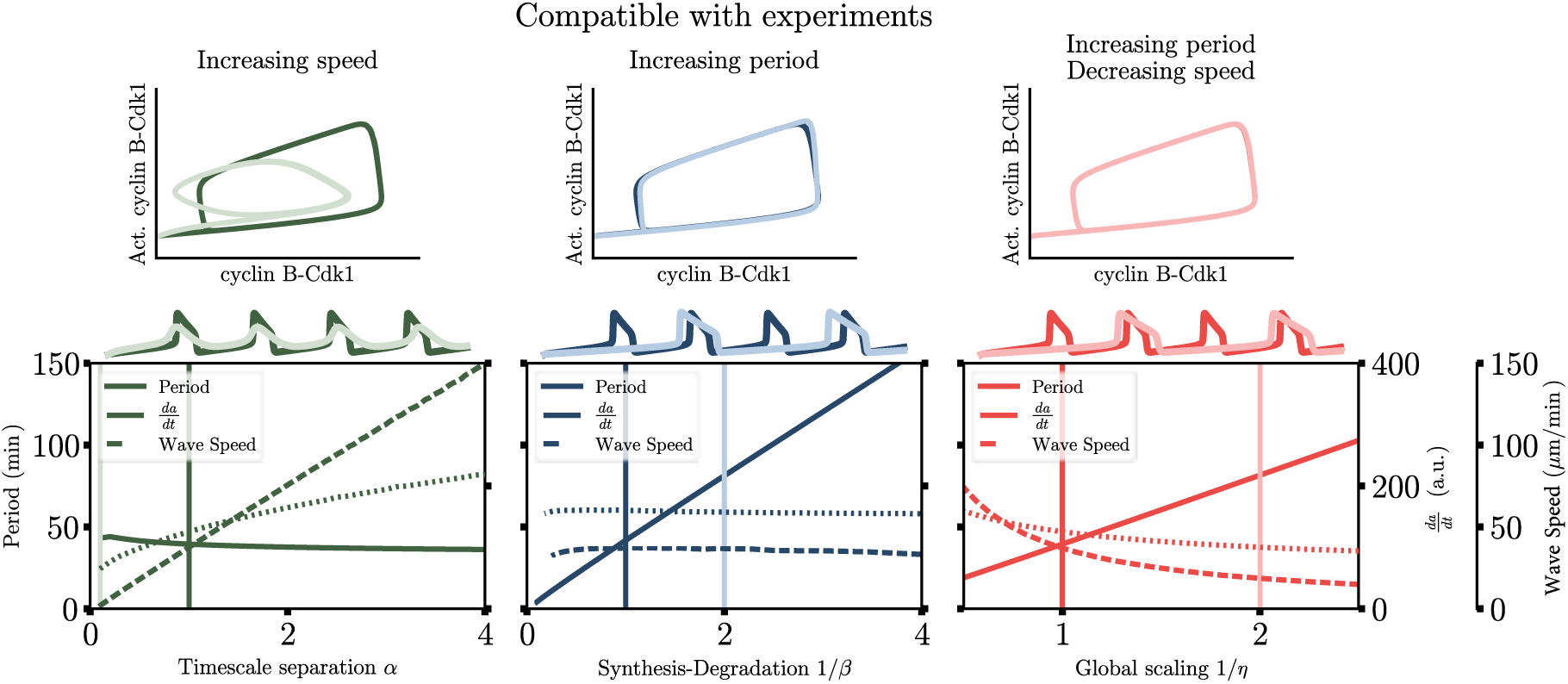
Dependence of the period, *da/dt*, and wave speed for different parameter scalings. Top: Phase space representation of the oscillator and the corresponding time series. Bottom: Period, *da/dt*, and wave speed is represented using continuous, dashed, and dotted lines as function of timescale separation *α* (left), decreasing synthesis and degradation rates with *β* (middle), and all rates with *η* (right). Vertical lines indicate the parameter values used in the top panels with the respective colors.

**Fig. S4.**
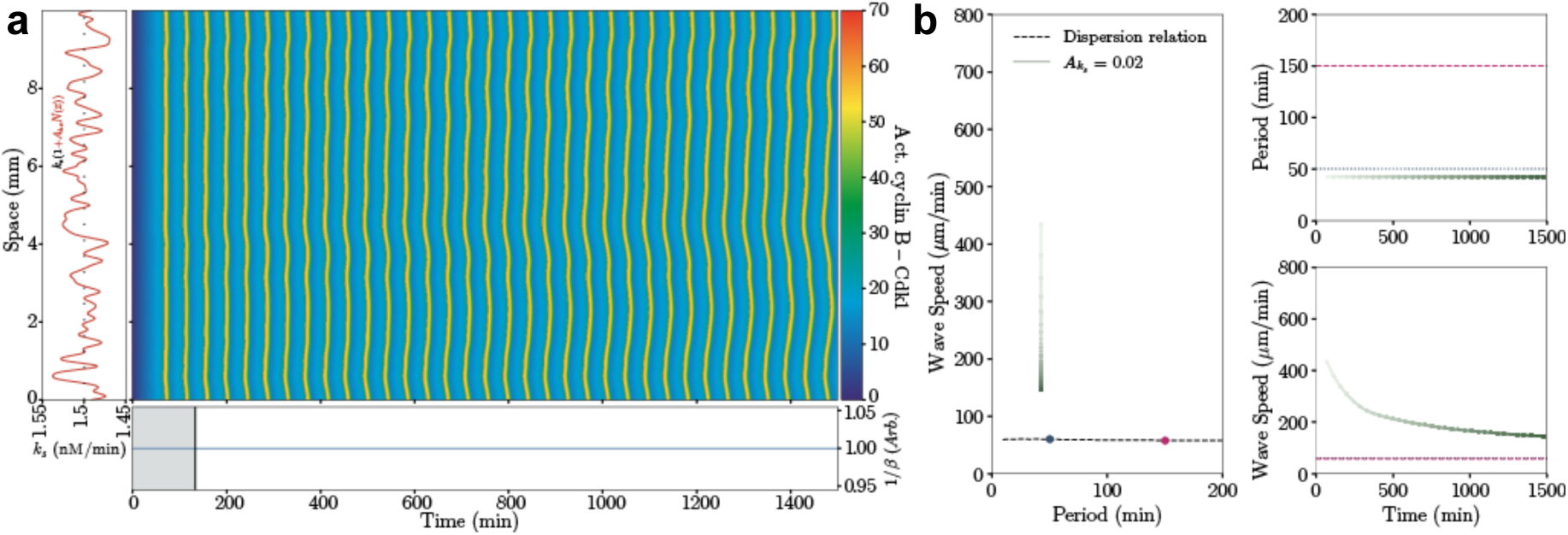
**a** Spatiotemporal evolution of the activity of cyclin B-Cdk1 showing the transition from phase to trigger waves with constant period with time using the parameter *β* (*t*) = 1 (bottom) and spatial variability in the synthesis term. **b** Speed-period relation and temporal dependence of the period and wave speed of the numerical simulation in panel **a**, and the theoretical dispersion relation using the parameter *β* to scan the period (black dashed line) same as in Fig. 2 included for comparison.

**Fig. S5.**
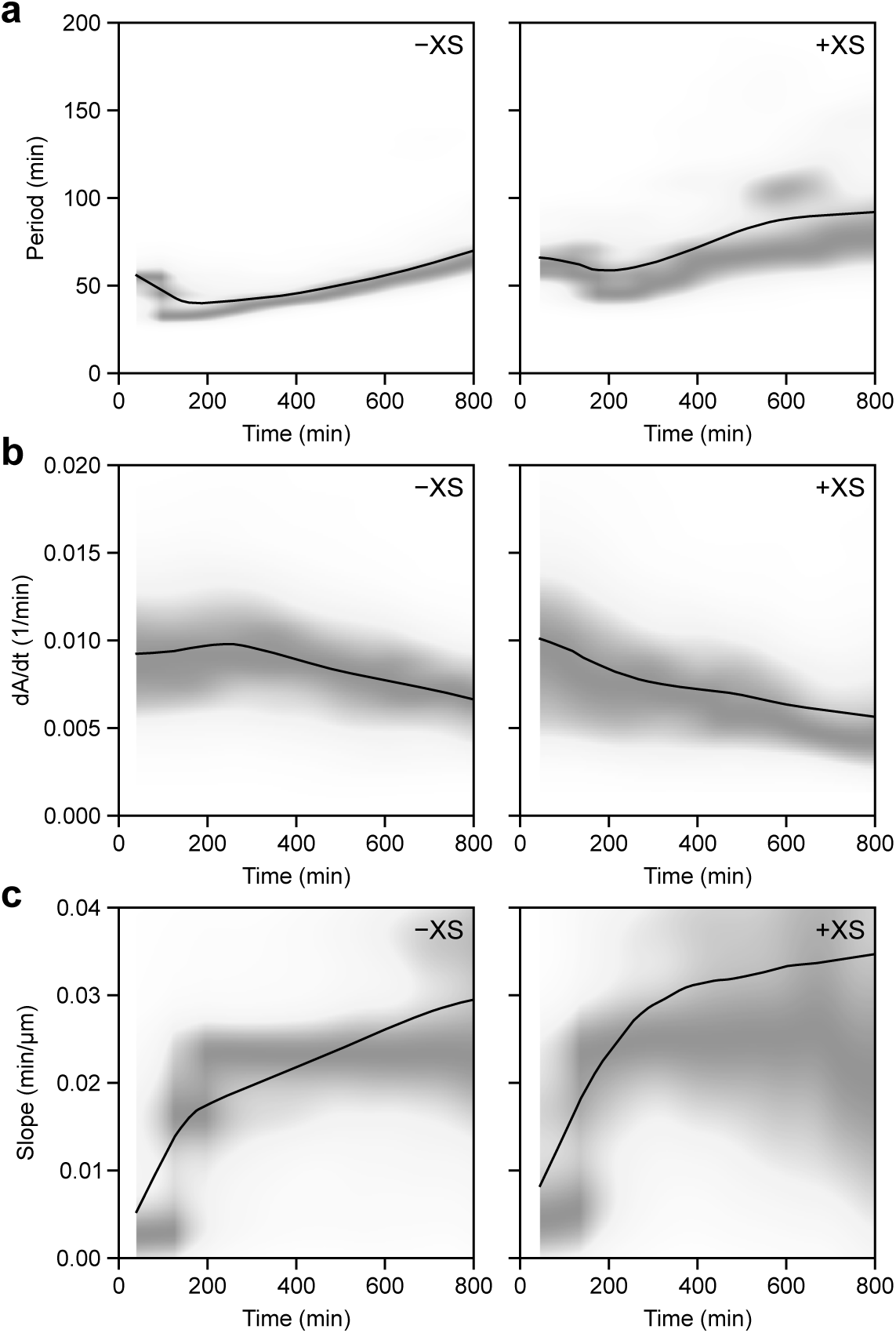
Period, maximum activation rate, and slope for CSF boundary-driven mitotic waves. **a** Period for no sperm DNA (*−*XS) and added sperm DNA (+XS) cases. Both columns feature the kernel density estimation over time with solid lines representing the LOWESS estimation. **b** Maximum activation rate. **c** Slope.

## MOVIES

**Mov. 1 Cdk1 wave dynamics in extracts without nuclei.** Data shared with the kymograph in Fig. 1. (Top) Pseudo-color movie of spatiotemporal dynamics of Cdk1 activity in bulk extracts. Color scale as in Fig. 1. Early times exhibit phase waves which give way to trigger waves over time. (Bottom) Detrended and smoothed FRET ratio (averaged over the width of the tube) plotted across the length of the tube. This shows how the spatial profiles develop from diffuse phase waves to pulse-like trigger waves.

**Mov. 2 Wave dynamics in extracts with reconstituted nuclei.** Data shared with the kymograph in Fig. 3. (Top) Pseudo-color movie of spatiotemporal dynamics of Cdk1 activity in bulk extracts. Color scale as in Fig. 3. Early times exhibit phase waves which quickly give way to trigger waves emanating from nuclei. Nuclei appear as hot-colored regions due to their import of active Cdk1. (Bottom) Detrended and smoothed FRET ratio (averaged over the width of the tube) plotted across the length of the tube. The curve represents the average over the width of the tube.

**Mov. 3 Excitable pulse in interphase extract driven by CSF.** Data shared with the kymograph in Fig. 4**b**. (Top) Pseudo-color movie of excitable pulse of Cdk1 activity in interphase extract as driven by CSF. (Bottom) Detrended and smoothed FRET ratio (averaged over the width of the tube) plotted across the length of the tube. A singular pulse is driven by the source.

**Mov. 4 CSF-driven wave dynamics in extracts without nuclei.**Data shared with the kymograph in Fig. 4**d**, left. (Top) Pseudo-color movie of trigger wave pulses in cycling extracts as driven by CSF. Phase wave dynamics are permanently abolished by driving. (Bottom) Detrended and smoothed FRET ratio (averaged over the width of the tube) plotted across the length of the tube. The curve represents the average over the width of the tube. The CSF source drives multiple trigger wave pulses.

**Mov. 5 CSF-driven wave dynamics in extracts with reconstituted nuclei.** Data shared with the kymograph in Fig. 4**d**, right. (Top) Pseudo-color movie of trigger wave pulses of Cdk1 activity in cycling extracts with reconstituted nuclei as driven by CSF. Both nuclei and the source drive trigger waves, but the CSF source ultimately dominates. (Bottom) Detrended and smoothed FRET ratio (averaged over the width of the tube) plotted across the length of the tube. The curve represents the average over the width of the tube.

